# *Skopi*: a simulation package for diffractive imaging of noncrystalline biomolecules

**DOI:** 10.1101/2021.12.09.471972

**Authors:** Ariana Peck, Hsing-Yin Chang, Antoine Dujardin, Deeban Ramalingam, Monarin Uervirojnangkoorn, Zhaoyou Wang, Adrian Mancuso, Frédéric Poitevin, Chun Hong Yoon

## Abstract

X-ray free electron lasers (XFEL) have the ability to produce ultra-bright femtosecond X-ray pulses for coherent diffraction imaging of biomolecules. While the development of methods and algorithms for macromolecular crystallography is now mature, XFEL experiments involving aerosolized or solvated biomolecular samples offer new challenges both in terms of experimental design and data processing. *Skopi* is a simulation package that can generate single-hit diffraction images for reconstruction algorithms, multi-hit diffraction images of aggregated particles for training machine learning classification tasks using labeled data, diffraction images of randomly distributed particles for fluctuation X-ray scattering (FXS) algorithms, and diffraction images of reference and target particles for holographic reconstruction algorithms. We envision *skopi* as a resource to aid the development of on-the-fly feedback during non-crystalline experiments at XFEL facilities, which will provide critical insights into biomolecular structure and function.

## 1. Introduction

The unique capabilities of X-ray free electron laser (XFEL) sources have led to significant advances in structural biology since the first hard X-ray laser began operation at the Linac Coherent Light Source (LCLS) in 2009 (Spence, 2018; Jamison, 2010). The XFEL technique of serial femtosecond crystallography (SFX) has proven particularly transformative for protein crystallographers. Ultra-bright X-rays enable studying crystals too small or radiation-sensitive for synchrotron sources, while the femtosecond pulses have permitted time-resolved studies of enzyme catalysis at atomic resolution (Schlichting, 2015; Stauch & Cherezov, 2018). By contrast, coherent X-ray diffractive imaging (CXDI) of non-crystalline biological samples has remained a largely nascent application, despite these experiments being the original intent of XFEL technology. Several studies have demonstrated successful single-particle imaging (SPI) of large targets such as viruses and organelles (Seibert *et al*., 2011; van der Schot *et al*., 2015; Brändén *et al*., 2019; Hantke *et al*., 2014), but 3D reconstruction has been limited to nanoscale resolution (Ekeberg *et al*., 2015; Rose *et al*., 2018; Kurta *et al*., 2017). Multi-particle techniques, including fluctuation X-ray scattering (FXS) (Mendez *et al*., 2014; Mendez *et al*., 2016; Kurta *et al*., 2017; Doniach, 2018; Pande *et al*., 2018) and holography (Gorkhover *et al*., 2018), have also been explored for structure determination. However, CXDI of noncrystalline biological samples remains far from routine, in contrast to the success of XFEL experiments of inorganic materials that provide significantly more signal (Ayyer *et al*., 2020). Improvements in resolution are needed before these techniques can address novel questions in structural biology.

The principal challenges faced by CXDI studies of noncrystalline samples are the low signal relative to instrumental background, sparsity of hits in the large volume of collected data, and intrinsic heterogeneity of the imaged particles (Bielecki *et al*., 2020; Daurer *et al*., 2017). Experimental advances have improved both sample delivery and the dynamic range of detectors, increasing the quality of the measured signal (Bielecki *et al*., 2019). Now some of the most pressing obstacles to achieving high-resolution reconstructions from these data are algorithmic in nature. In particular, rigorous classification is required both to identify the useful single-particle hits from shots of aggregates or no particles and to cluster images based on conformational heterogeneity (Maia *et al*., 2009; Yoon *et al*., 2011; Reddy *et al*., 2017; Cruz-Chú *et al*., 2021). In addition, more sophisticated background subtraction methods are needed to isolate the low-intensity particle scattering prior to reconstruction. While methods to perform classification and background removal have been explored and applied to the few experimental datasets available (Assalauova *et al*., 2020; Shi *et al*., 2019), it is unclear how well these algorithms will generalize to new experiments.

Realistic simulations of CXDI experiments could accelerate the maturation of this field by providing a test-bed for advanced data pre-processing and reconstruction algorithms. *Condor* is another open-source software available for CXDI simulations and was created to facilitate planning experiments at first-generation XFELs (Hantke *et al*., 2016). However, the ongoing development of different types of single particle imaging experiments with higher repetition rates and more sophisticated detectors poses new challenges and thus calls for tools capable of simulating these technological advances at scale. Here we present *skopi*, a software package designed for rapid and high-throughput simulations of SPI, FXS, and holography experiments on modern detectors. The simulated experiments are highly customizable and readily support modeling the beam characteristics and detectors available at LCLS. *Skopi* also makes it convenient to include diverse sources of noise, including fluctuating dark noise, beam miscentering, a static sloped background, and fluence jitter, all of which can impact reconstruction. Additional challenges inherent to biomolecular imaging, such as variable particle hydration and heterogeneity, can be simulated as well. We anticipate *skopi* will be a useful resource for developing new data processing algorithms, guiding experimental design, and in time aiding on-the-fly feedback during CXDI experiments of noncrystalline biological samples.

## 2. Modular implementation

The design of *skopi* is highly modular, with the three main components of a CXDI experiment — the particle, beam, and detector — agnostic to the type of experiment being simulated (Fig. 1). Each component can be modeled with a range of complexity, from the ideal case to more sophisticated representations that mimic experimental errors and noise. We briefly describe these components below.

**Figure 1.**
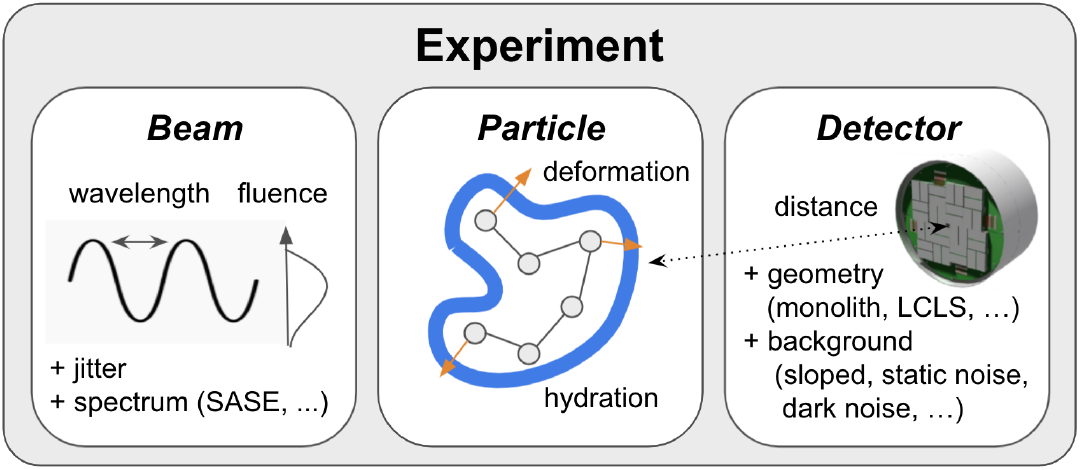
Modular architecture. The three principal components of each experiment – the beam, particle, and detector – are initialized independently of each other. Once these components are set up, diffraction patterns from a range of CXDI experiments can be efficiently simulated.

The particle object stores the biomolecule’s atomic coordinates and associated form factors. Conformational heterogeneity can be modeled by sampling along the particle’s normal modes to generate alternate states; this is accomplished using the anisotropic network model from the *ProDy* package (Bakan *et al*., 2011) (Fig. S1A). Another source of sample heterogeneity is the disordered solvent shell surrounding aerosolized particles, which helps to preserve structural integrity in vacuum but increases shot-to-shot variation (Hau-Riege *et al*., 2007; Mandl *et al*., 2020). To account for this effect, *skopi* enables modeling a hydration layer that follows the particle’s contour and has a variable width between shots (Fig. S1B).

The beam object contains information about the beam’s dimensions, fluence, and wavelength spectrum. The simplest case models a single monochromatic spike with spatially uniform fluence. A more advanced option accounts for self-amplified spontaneous emission (SASE), which produces ultrabright pulses but broadens the beam’s energy spectrum (Geloni *et al*., 2017). *Skopi* models SASE spectra as trains of uncorrelated spikes using a Gaussian kernel density estimate to approximate their energy distribution (Fig. S2). Noise can be added by modeling spatial variation of the fluence in the plane of the beam and by introducing jitter, in which shot-to-shot changes in total fluence are Gaussian-distributed.

The simplest detector is a monolithic square pixel array of user-defined dimensions. The diverse detectors in use at LCLS for CXDI experiments are also supported (Fig. S3A). Recently developed detectors, such as the Jungfrau and ePix10k, are implemented with an auto-ranging feature that avoids saturation to increase their dynamic range (Blaj *et al*., 2015; van Driel *et al*., 2020; Redford *et al*., 2018) (Fig. S3B). LCLS detectors also have access to the calibration constants and fluctuating dark noise from specific experiments (Fig. S4A). Once initialized with a beam, the detector object is populated with information about the pixels’ position in reciprocal space, solid angle distended, and polarization. Finally, the detector object also supports modeling both beam miscentering, in which the beam’s focus is displaced relative to the detector’s center, and a user-defined sloped background to model parasitic scattering, which is a source of correlated noise that cannot be averaged away by merging more data (Fig. S4B-C).

Once the particle, beam, and detector are set up, any of the three principal CXDI experiments can be easily and efficiently simulated. *Skopi* provides convenient interfaces for each of the SPI, FXS, and holography experiment types, from which small or wide angle X-ray scattering (SAXS or WAXS) profiles can be derived through radial averaging. These interfaces act as wrappers for a common experiment class, recognizing that CXDI experiments are at their core very similar, differing only in the number, type, and relative position(s) of particle(s) in the beam (Fig. 2). For each shot, the particle or set of particles is randomly displaced and oriented in the volume intersected by the sample delivery jet and beam, and the coherent diffraction pattern is computed. To facilitate incremental testing of reconstruction algorithms, either the ideal intensities or quantized photons can be saved, in addition to the number of particles, particle orientations, and particle positions at each shot. Beyond Poisson error, more sophisticated types of noise can also be defined in configuring the experiment and individually tuned to achieve the desired signal-to-noise ratio.

**Figure 2.**
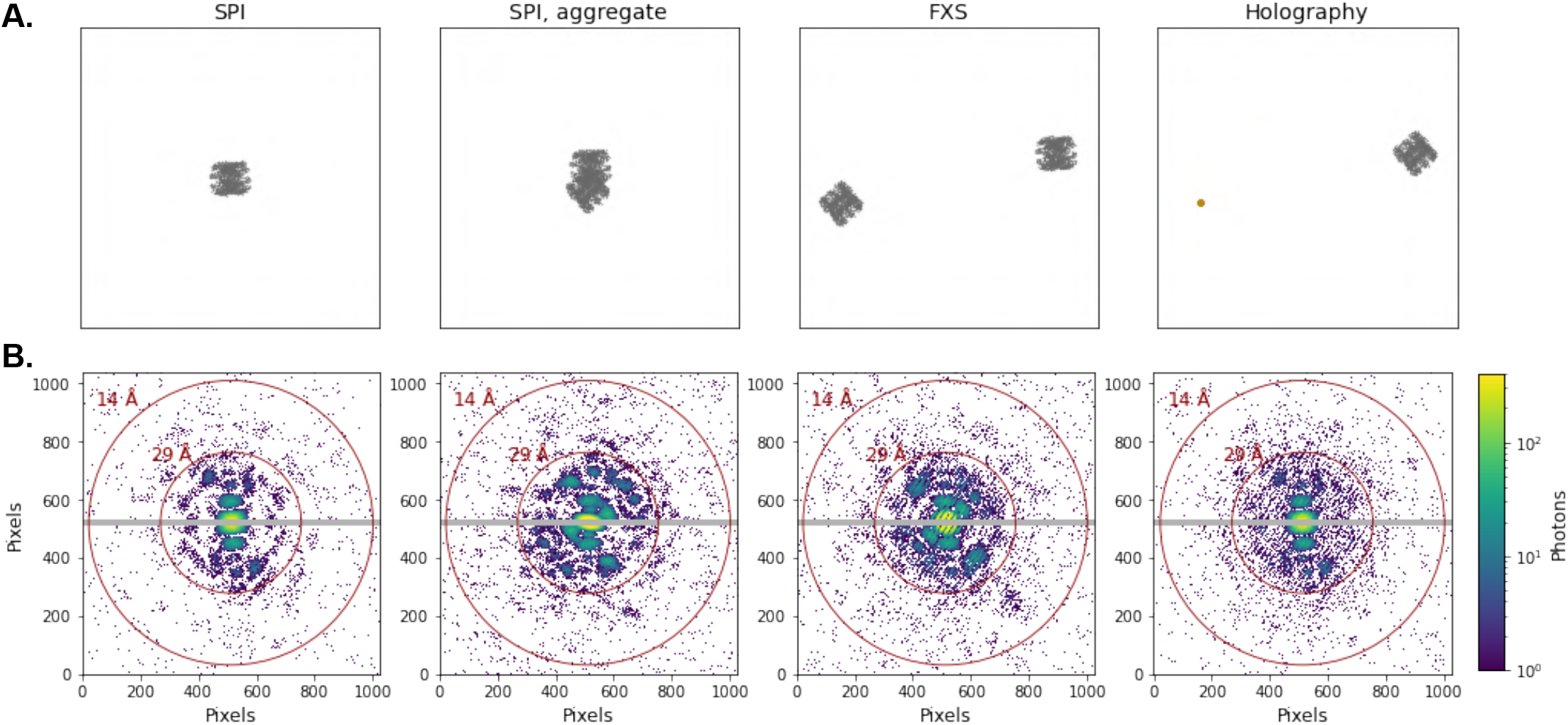
Overview of noncrystalline CXDI experiments. *Skopi* supports simulations of SPI, FXS, and holography experiments. (A) The projection of the particle(s) in the plane of the beam and (B) the corresponding diffraction patterns are shown for each experiment type. In the case of SPI, either an individual particle or an aggregate can be simulated. In FXS and holography experiments, multiple particles are in the beam; for holography, one of these particles serves as a reference – in this case, a small cluster of gold atoms. The biomolecule used in these simulations is a chaperonin (PDB ID: 3iyf). In (B) the grey region marks the gap between panels of the PnCCD detector. The beam fluence was artificially inflated to aid visualization.

## 3. Benchmarking

*Skopi* achieves rapid and high-throughput simulations of CXDI experiments through graphical processing unit (GPU) acceleration and parallelization. Because diffraction calculations scale linearly with particle size, the full 3D diffraction volume for each particle is computed when an experiment is first configured and thus represents a one-time cost (Fig. S5A). Diffraction images are then efficiently calculated by slicing through this pre-computed volume (Fig. S5B). The disadvantage of this approach is the inability to account for the stochastic effects of radiation damage during each interaction between the sample and beam; however, radiation damage is anticipated to be minimal for biological samples, particularly in comparison to instrumental background (Neutze *et al*., 2000; Östlin *et al*., 2019; Spence, 2017). On a single NVIDIA GeForce GPU, an SPI dataset acquired on a one megapixel PnCCD detector could be simulated at a rate of 0.6 diffraction patterns per second. This rate was reduced to 0.4 images per second for the SPI aggregate, FXS, and holography experiments due to the need to distribute clustered or dispersed particles in the beam for each shot. Dataset generation can be easily parallelized across an arbitrary number of GPU nodes to increase throughput.

## 4. Validation

We validated *skopi* by recovering the protein structure from simulated diffraction images of a chaperonin (PDB ID: 3iyf). SPI datasets consisting of 5k images each were generated in the absence or presence of noise on a PnCCD detector positioned for a resolution limit of 14 Å at the edge of the detector. Reconstruction was performed using a Cartesian implementation of the multi-tiered iterative phasing (MTIP) algorithm (Donatelli *et al*., 2017; Dujardin *et al*., 2020). As expected, the protein structure was accurately recovered to a resolution of 15 Å from noise-free diffraction images (Fig. 3). Poisson noise reduced the resolution of the recovered structure to 20 Å, and additionally introducing beam jitter degraded reconstruction quality further, as evident in the loss of the protein’s eight-fold symmetry (Fig. 3). *Skopi* thus provides a useful tool to assess the tolerance of reconstruction algorithms to different types of noise.

**Figure 3.**
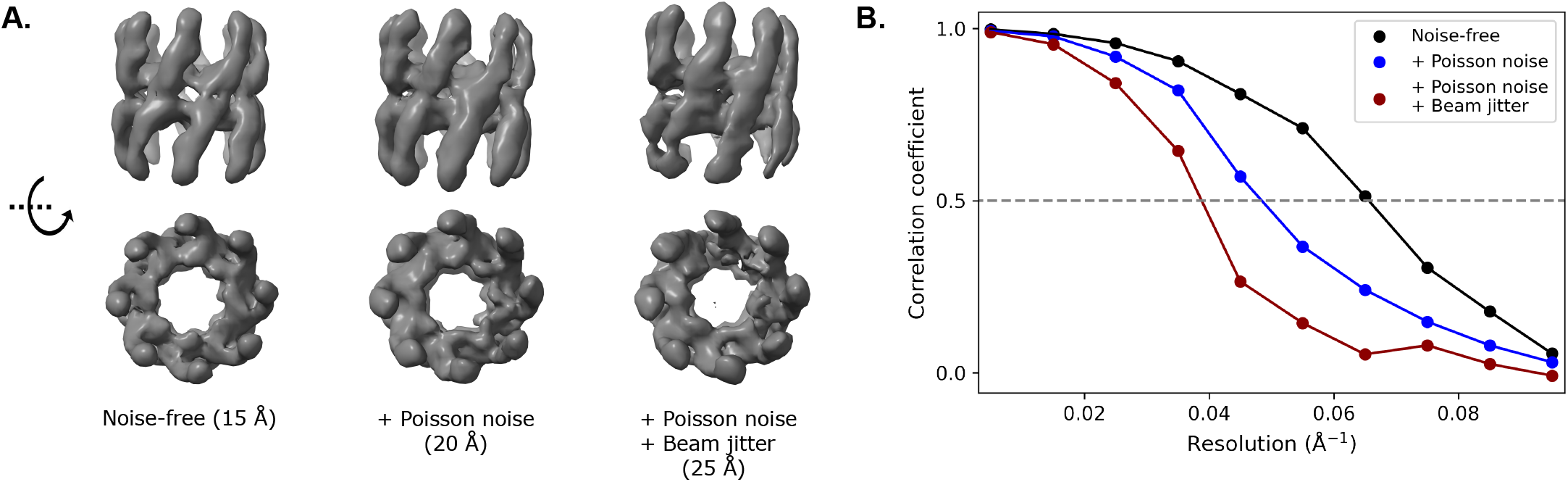
Reconstruction from simulated SPI datasets. SPI datasets of a chaperonin were simulated in the absence or presence of noise. A multi-tiered iterative phasing algorithm was used to recover the protein structure from 5k images of the indicated dataset. (A) Isosurfaces of the density map reveal the loss of eight-fold symmetry with increasing noise. The resolution of each reconstruction is noted in parenthesis. (B) The resolution was measured as the spatial frequency at which the Fourier shell correlation (FSC) between the reconstructed and reference maps dropped to 0.5.

## 5. Conclusion

Here we present *skopi* as a convenient tool to efficiently simulate CXDI experiments of noncrystalline biological samples. The modular design of this package makes it easy to represent each component of an XFEL experiment with a range of complexity, including modeling the most up-to-date features of current LCLS detectors (van Driel *et al*., 2020). Another focus of *skopi* is to produce simulated data with realistic noise. The different sources of error that typify CXDI experiments — including fluence jitter, beam miscentering, and fluctuating dark noise — can be readily incorporated into the simulation without additional scripting.

The paucity of experimental CXDI datasets of noncrystalline biological samples has hindered algorithmic growth in the field. We anticipate that *skopi* could help fill this gap by its ability to rapidly provide diverse simulated datasets with realistic noise. This may prove particularly valuable for the development of machine learning algorithms to perform classification, since labeled data can easily be generated in bulk. Classification algorithms would not only benefit data pre-processing to separate single-particle hits from aggregate shots, but could also be used during reconstruction to sort diffraction images based on the particle’s conformation (Ignatenko *et al*., 2021), as done in cryo-electron microscopy (Punjani & Fleet, 2021; Zhong *et al*., 2021; Chen & Ludtke, 2021).

In addition to aiding algorithm development, we envision *skopi* as a critical tool for guiding experimental data collection at LCLS and other XFEL facilities through large-scale start-to-end facility simulations (Yoon *et al*., 2016; Fortmann-Grote *et al*., 2017). By supporting both the advanced detectors currently in use at LCLS and the inclusion of fluctuating dark noise from past experiments, *skopi* enables simulating highly realistic diffraction patterns that would be obtained at this facility. These simulated datasets in turn provide an estimate of the available signal under different experimental conditions, potentially allowing settings to be optimized and offering an estimate of the volume of data required for reconstruction in advance of XFEL experiments.

## Appendix A Theory, Implementation and Usage

### Elements of theory

In this section, we summarize the main elements of coherent X-ray optics taken from (Paganin, 2006) that allow us to derive the diffraction model implemented in *skopi*. Propagation of electromagnetic waves Ψ(**r**, *t*) in nonmagnetic matter follows the wave equation, where the local velocity is modulated by the local electric permittivity *ϵ* while the magnetic permeability *μ*_0_ is assumed to be that of free space everywhere:

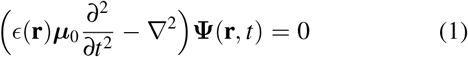

Separation of temporal and spatial variables is assumed through spectral decomposition:

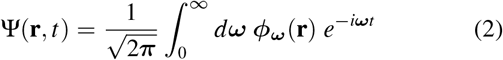

where the frequency *ω* of each monochromatic wave is related to its wavevector modulus *k* through the velocity of light in free space: *ω* = *kc*. The effect of local perturbations can be factored in the refractive index 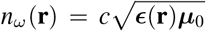, which measures the relative velocity of light in the medium compared to free space. Inserting the spectral decomposition of the electromagnetic wave in the wave equation yields an inhomogeneous Helmholtz equation for each monochromatic wave:

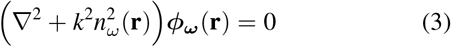

In free space, Eq.3 simplifies to the homogeneous Helmholtz equation which can be solved and yields the following solution to the wave equation 1: Ψ(**r**, *t*) = exp *i*(**k**.**r** - ***ω**t*). The spatial wavelength λ of this wave is such that cos *k*λ = 1, so the wavevector modulus is related to it through the relation 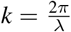.

More generally, it can be shown that Eq. 2 is equivalent to its integral-equation formulation, which involves the incident wave *ϕ*^(0)^ and a convolution with the Green’s function 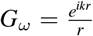:

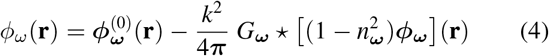

In the Born approximation, the wave function on the righthand side of Eq. 4 is assumed to be identical to the incident wave function, thus trivializing the integral equation formulation. Further assuming the case of Fraunhofer (far-field) diffraction and of an incident plane wave 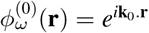, we can introduce the scattering amplitude *f* (Δ**k**) where 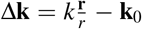:

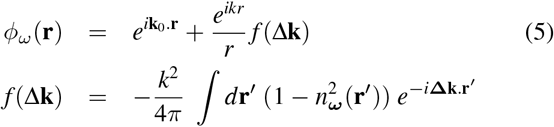

Assuming a specific relationship between the refractive index and the electron density of the object *ρ*, the scattering amplitude is scaled by the Thomson scattering length *r*_0_:

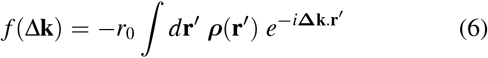

Finally, noting 2*θ* as the angle between the incident and resulting wave vectors, we introduce the notation **s** = **Δk** where 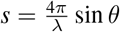. In the superposition approximation, the molecular electron density is approximated as a sum of individual atomic contributions. In diffraction space, the atomic form factors *f_i_* are tabulated as a function of sin *θ*/λ:

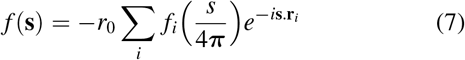

The resulting structure factors *f* can then be corrected for beam polarization and solid angle effects and corrupted with several types of noise that we encompass into functions *α* and *β* to yield the intensity *I*. The diffraction intensity is quantized by drawing from the corresponding Poisson distribution 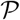 to simulate the photon counts *C* measured on the detector:

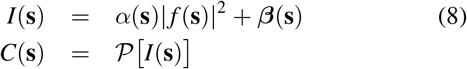

### Implementation

In practice, Eq. 7 is pre-computed on a discrete 3D cubic grid whose reciprocal extent is determined by the detector and beam parameters. Once the resulting diffraction volume is available in memory, it can readily be sliced at an arbitrary orientation and curvature in two steps. First, the detector pixels are mapped to their corresponding positions in the reciprocal volume. Second, the complex structure factor at each pixel is estimated through trilinear interpolation. The resulting diffraction intensities are then adjusted to account for beam polarization, solid angle distortion, and diverse sources of noise.

### Usage

The Python script below illustrates how the modularity of *skopi* is implemented in practice. Classes for the beam, particle and detector are instantiated and given to the experiment class, which in turn generates diffraction patterns.

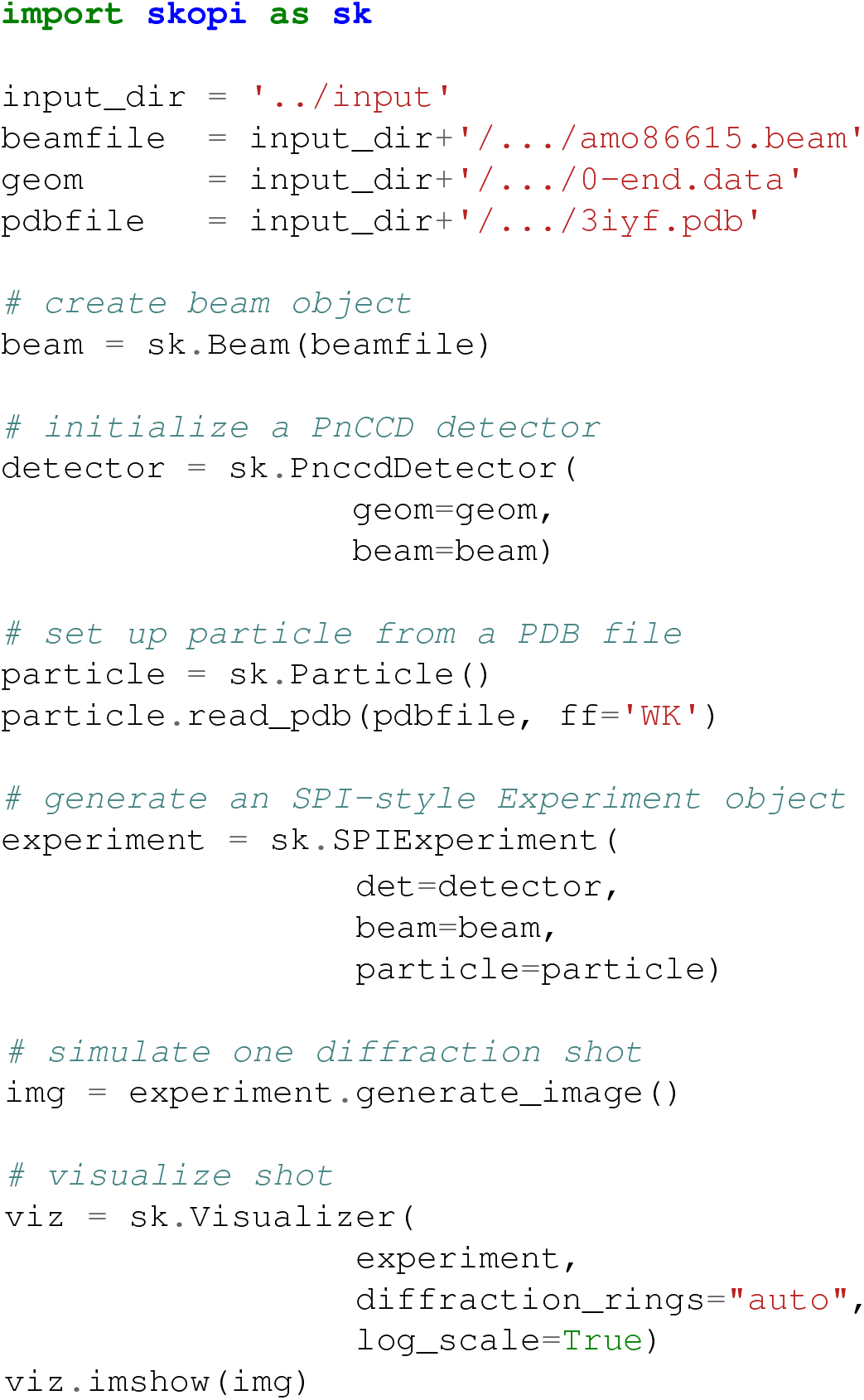

## Acknowledgements

We thank Haoyuan Li and Juncheng E for their contributions to the code. This research was supported by the Exascale Computing Project (17-SC-20-SC), a collaborative effort of the U.S. Department of Energy Office of Science and the National Nuclear Security Administration. Use of the Linac Coherent Light Source (LCLS), SLAC National Accelerator Laboratory, is supported by the U.S. Department of Energy, Office of Science, Office of Basic Energy Sciences under Contract No. DE-AC02-76SF00515.

---

**Figure S1.**
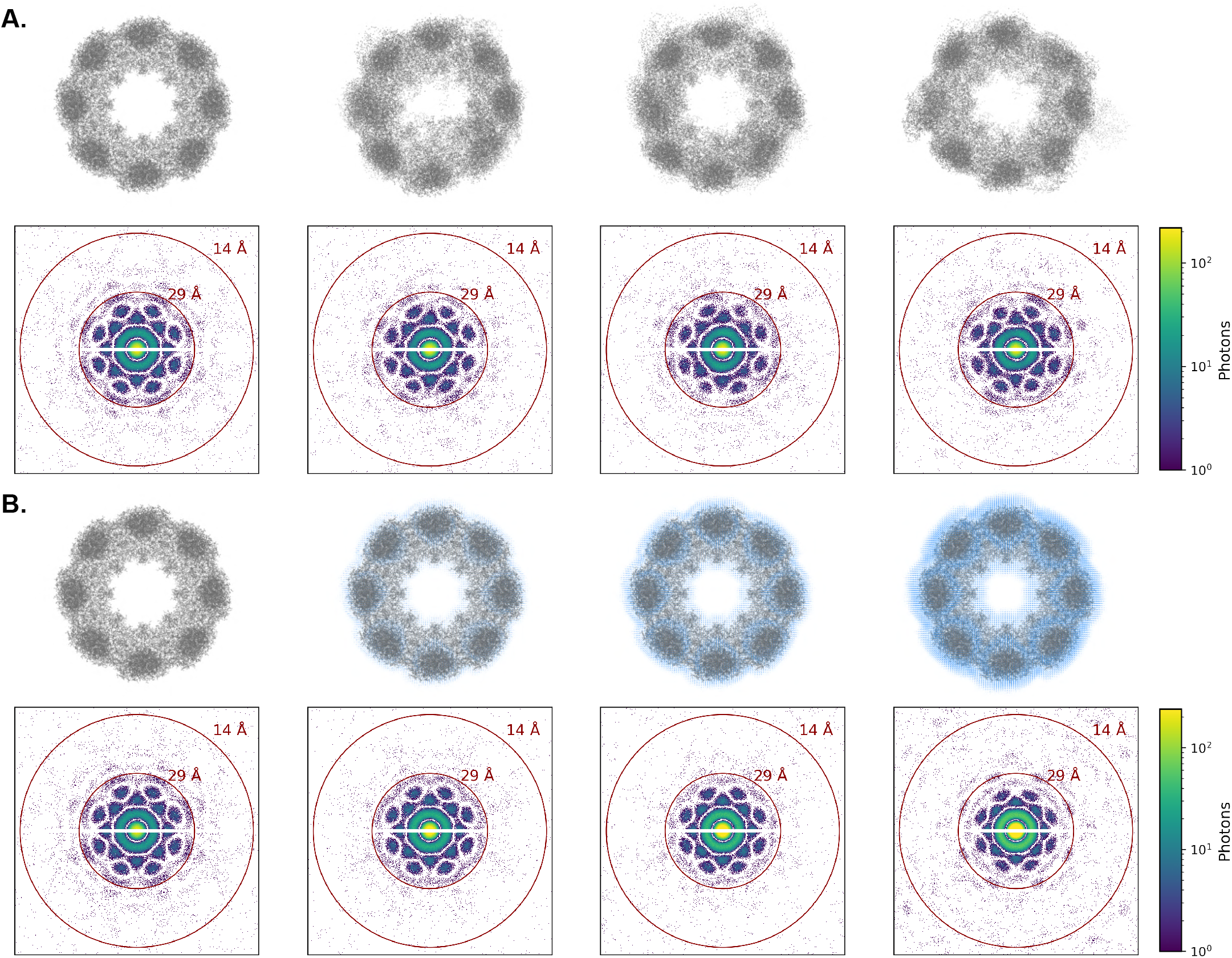
Sources of sample heterogeneity. (A) Distinct conformations were generated by sampling along the protein’s normal modes (*upper*), and the corresponding diffraction patterns are visualized (*lower*). Conformational heterogeneity results in visible distortion of the protein’s eight-fold symmetry along this axis in the simulated shots. (B) The hydration layer surrounding aerosolized particles is modeled as a disordered solvent shell of variable width that follows the contours of the protein. From left to right, the chaperonin protein (grey, PDB ID: 3iyf) was encased in a solvent shell (blue) of 0, 4, 8, and 12 Å (*upper*). The diffraction pattern for each solvation state is shown (*lower*).

**Figure S2.**
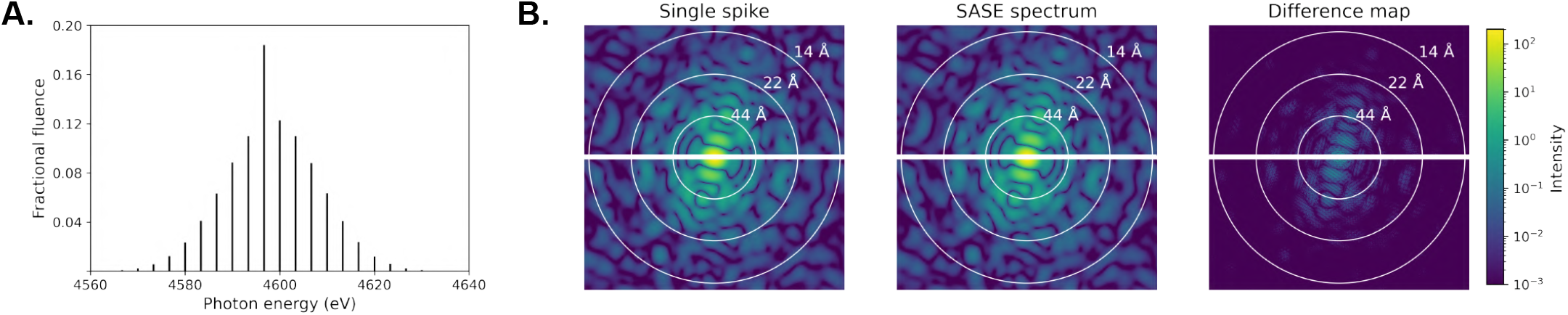
Modeling the beam as a SASE spectrum. (A) The distribution of spike energies and fractional fluences in a representative SASE spectrum are shown for a mean beam energy of 4600 eV. (B) Diffraction patterns for a particle in the same orientation but imaged with either a single, monochromatic spike or a SASE-style beam are compared, with the absolute difference map shown on the right. Because the magnitude of differences is on the same scale as quantization error, intensities rather than photons were computed. However, the source of the difference does not follow a Poisson distribution.

**Figure S3.**
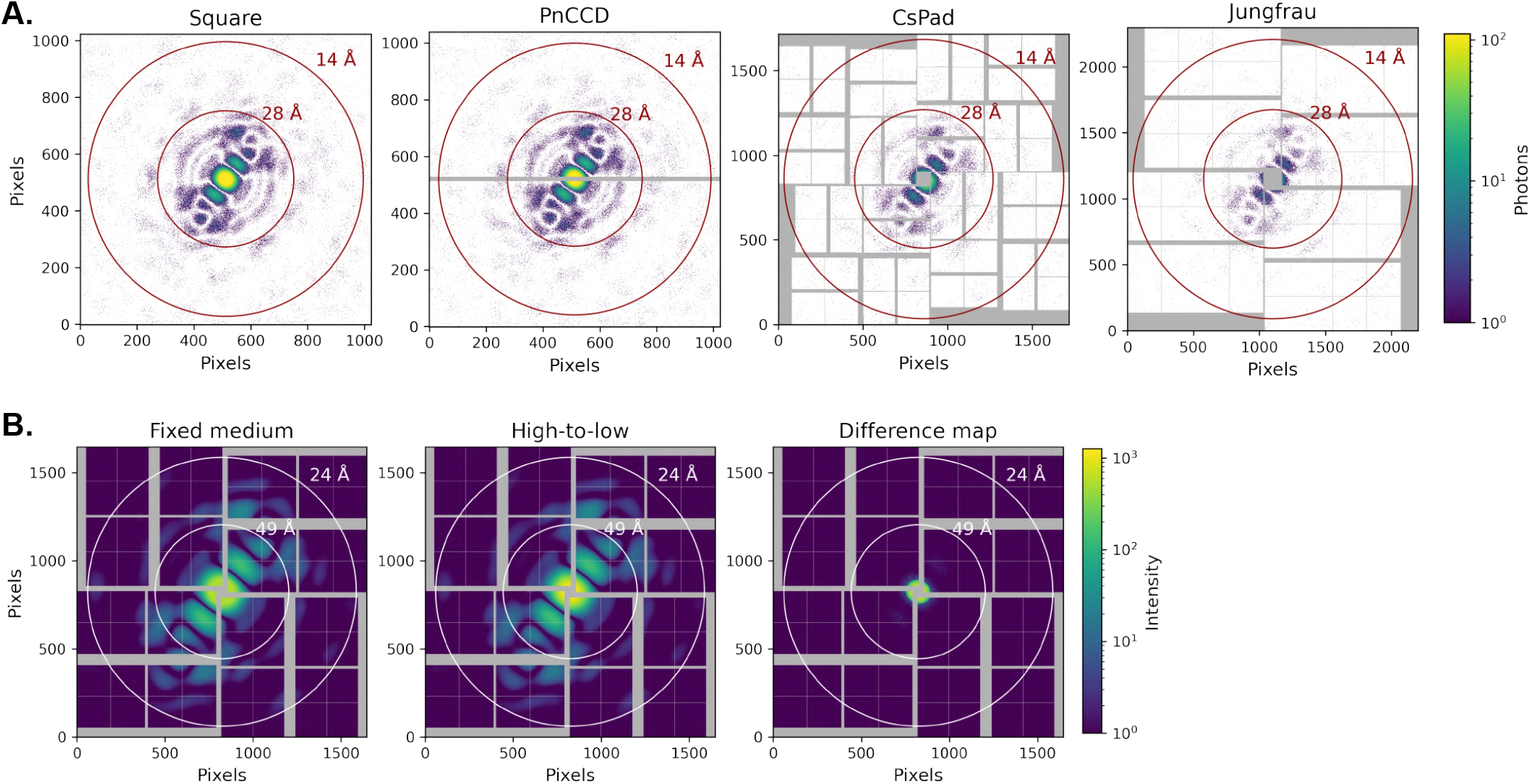
Support of diverse detector types. (A) In addition to a user-defined monolithic square detector, *skopi* supports simulating images on a range of LCLS detectors such as the PnCCD, CsPad, and Jungfrau. Diffraction patterns using the same beam and particle objects were simulated on each of these detectors. (B) The recently-developed Jungfrau and Epix10k detectors have an auto-ranging feature with various gain combinations to increase the dynamic range. Diffraction patterns simulated on the Epix10k are compared for the fixed medium and high-to-low gain combinations, showing that the latter mode eliminates saturation at high intensity values. Intensities rather than photons are shown and the detector distance has been adjusted to better illustrate where differences between the gain combination modes are most pronounced.

**Figure S4.**
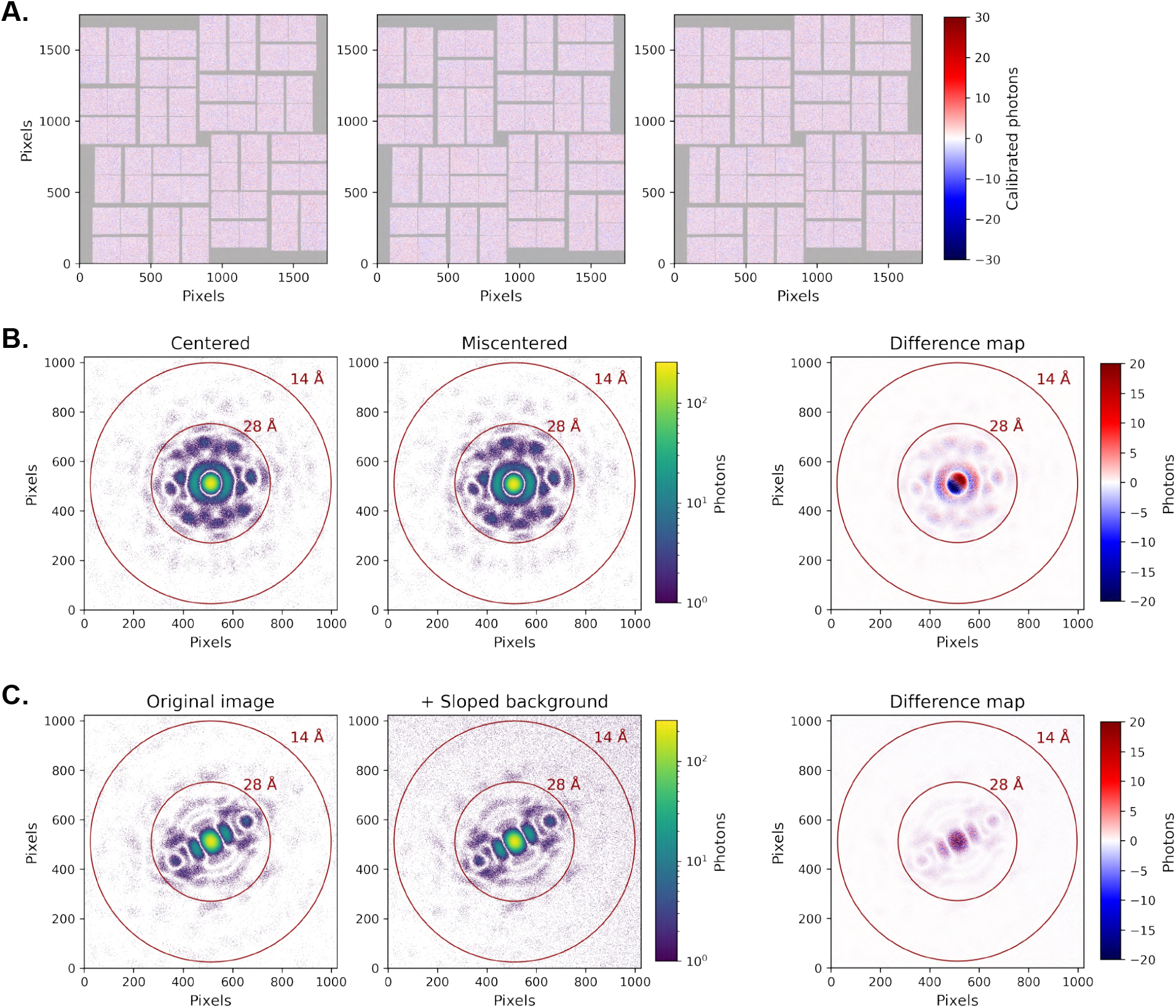
Sources of instrument noise. (A) Representative dark shots are shown from an experiment performed at LCLS on a CsPad detector. The dark noise fluctuates between images, and the counts can be negative due to pedestal subtraction. (B) Beam miscentering and (C) a user-defined sloped background can be introduced as additional sources of noise. Difference maps are shown on a linear scale to aid visual comparison.

**Figure S5.**
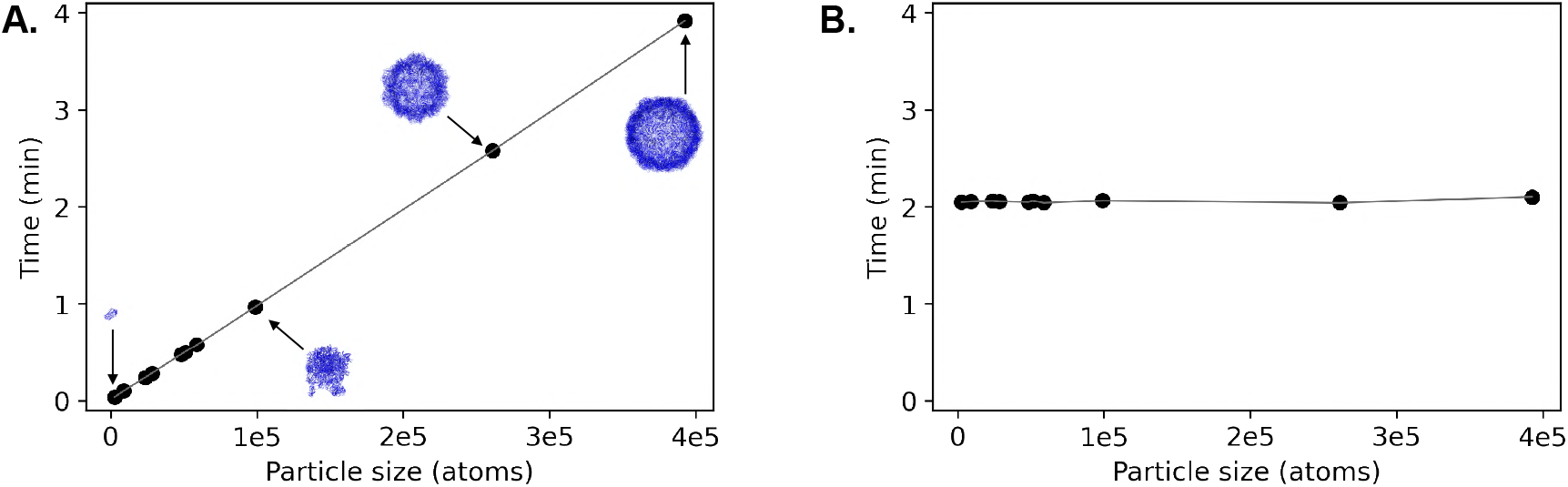
Scaling of the diffraction calculation with particle size. (A) The time required to initialize a (non-aggregate) SPI experiment is compared for different particle sizes and a square detector of roughly a quarter million pixels. Most of the elapsed time is spent computing the 3D diffraction volume, which scales linearly with particle size and is performed on a single GPU. Models of select particles are shown, from largest to smallest: rhinovirus (PDB ID: 2rmu), feline panleukopenia virus (PDB ID: 1fpv), 50S ribosomal subunit (PDB ID: 3cc4), and sialic acid binding protein (PDB ID: 2cex). (B) The elapsed time to simulate 100 diffraction patterns on a single GPU across a range of particle sizes. Once the experiment is set up, the diffraction image computation time is constant.

